# Brainstem circuit for sickness-induced sleep

**DOI:** 10.1101/2025.03.09.642181

**Authors:** Dana Darmohray, Yuanyuan Yao, Jiao Sima, Chien-Hao Chen, Daniel Silverman, Changwan Chen, Yang Dan

## Abstract

Increased sleep induced by immune activation plays a crucial role in facilitating recovery from illness. However, the neural mechanisms underlying sickness-induced sleep remain poorly understood. Here, we identify a brainstem circuit originating in the nucleus of the solitary tract (NST) that mediates sickness-induced sleep. Using activity-dependent genetic labeling, we tagged NST neurons activated by lipopolysaccharide (LPS) injection and showed that their chemogenetic activation strongly promotes non-rapid eye movement (NREM) sleep. These NST neurons project extensively to the parabrachial nucleus (PB), where LPS-activated neurons also promote NREM sleep. Fiber photometry imaging of several wake-promoting neuromodulators using their biosensors showed that evoked norepinephrine (NE) release from locus coeruleus (LC) neurons is markedly reduced by either LPS injection or direct activation of NST or PB sickness neurons. These results suggest that sickness-induced sleep is mediated in part by a brainstem circuit that regulates neuromodulator signaling.

## INTRODUCTION

Sleep is a highly conserved innate behavior that is indispensable for health and survival. It supports various cognitive and physiological processes, including memory consolidation, emotional processing, waste clearance, and metabolic regulation ^1–6^. Sleep also interacts bidirectionally with the immune system ^7–11^. Sleep loss leads to immune system dysregulation and ultimately death^12–14^. Conversely, following an immune challenge, the amount of time the animal spends in sleep increases substantially ^15,16^. Such an increase in sleep promotes functional recovery and survival during sickness or injury ^7,17,18^. However, the mechanisms by which the immune system regulates sleep are not well understood.

A range of sickness-induced behavioral changes, collectively known as sickness behavior, can be triggered by cytokines – signaling molecules that play crucial roles in regulating immune responses ^19–22^. Cytokine signals in the periphery can reach the brain through neural and humoral routes. The neural pathway begins with the sensory division of the vagus nerve, which projects to the nucleus of the solitary tract (NST) in the dorsal medulla. Vagotomy or reversible inactivation of the NST strongly abrogates sickness behavior, indicating a crucial role of this pathway in mediating the immune-brain communication ^23–27^. Recent studies have yielded important insights into how this pathway controls sickness behavior by identifying the subsets of vagal axons that sense peripheral inflammation and the NST neurons that drive sickness behavior ^24,28^.

While these studies elucidate the neural entry point for sickness behavior in general, the downstream pathways mediating specific symptoms such as increased sleep remain unclear. The NST is widely interconnected with brain areas that regulate functions spanning autonomic outflow to motivated behavior ^29–35^, allowing it to orchestrate multiple physiological and behavioral responses to sickness. Here, we explore the neural pathways from the NST that drive sleep during peripheral immune activation. Using activity-dependent genetic labeling and chemogenetic manipulation, we show that sickness-activated NST neurons and their projection target – the parabrachial nucleus (PB) – can promote sleep in the absence of inflammation. Using genetically encoded fluorescence-based GRAB sensors for several wake-promoting neuromodulators, we show that evoked norepinephrine (NE) release from the locus coeruleus (LC) is markedly reduced by peripheral inflammation or direct activation of NST or PB sickness neurons. These results suggest that sickness-induced sleep could be mediated in part by a brainstem circuit that regulates neuromodulator signaling.

## RESULTS

### NST sickness-activated neurons promote sleep

To elucidate the neural circuitry underlying sickness-induced sleep, we first tested whether stimulating sickness-activated NST neurons is sufficient to increase sleep in the absence of peripheral immune activation. We used activity-dependent genetic labeling ^36^ to tag sickness-activated neurons in the NST. TRAP2 mice expressing an inducible Cre (2A-iCreER^T2^) under the *Fos* promoter were crossed to a reporter line expressing eGFP (Fig. 1A). Lipopolysaccharide (LPS; 0.4 mg/kg) was injected intraperitoneally (IP) to elicit sickness behavior together with tamoxifen (4-OHT; 20 mg/kg). After >7 days, we found significantly more eGFP-labeled neurons in the NST (referred to as “NST^LPS-TRAP^” neurons) compared to saline-injected control mice (Fig. 1A, Fig. S1A; right; *t*-test, *t* = -8.1, *p* = 0.00002).

**Figure 1.**
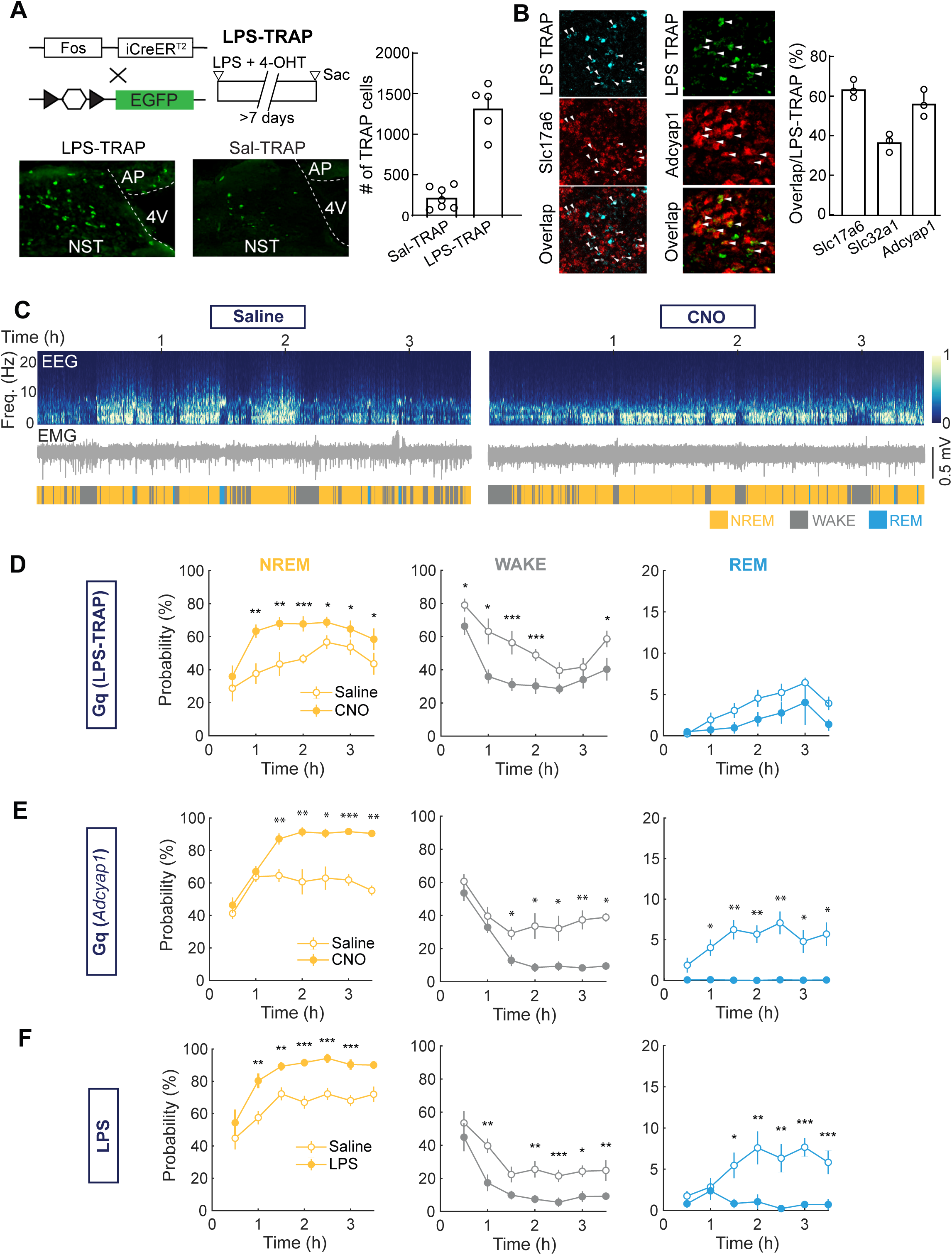
NST sickness-activated neurons promote sleep. **(A)** *Top left*: Schematic of experimental protocol. Mice expressing inducible Cre (2A-iCreER^T2^) under the *Fos* promoter (TRAP2) were crossed with Cre-inducible reporter mice expressing eGFP. LPS (0.4 mg/kg) and tamoxifen (4-OHT; 20 mg/kg) were co-injected to label sickness-responsive neurons (LPS-TRAP). *Bottom left*: Example of eGFP labelled NST neurons in LPS-TRAP compared with Sal-TRAP (tamoxifen and saline co-injection) mice. *Right:* Average number (±SEM) of TRAP NST neurons (eGFP labeled) in saline (SAL-TRAP, n = 7) and LPS conditions (LPS-TRAP, n = 5). Open circles represent individual mice. **(B)** Average overlap (±SEM) between LPS-TRAP neurons with cell type markers, *Slc17a6* (glutamatergic), *Slc32a1* (GABAergic/glycinergic), and *Adcyap1* shown by double FISH (*left*) and overlap quantification (*right*). Open circles represent individual samples. **(C)** Example experiment for chemogenetic activation of NST^LPS-TRAP^ neurons (Gq (LPS-TRAP)). EEG, EMG, and color-coded brain states are shown over 3-h recording session for Saline (*left*) and CNO injected mouse (*right*). **(D)** Average changes in NREM, wake and REM following Saline (open circles) or CNO injections (solid circles, n = 9) in mice expressing Gq DREADD in NST^LPS-TRAP^ neurons (Gq (LPS-TRAP)). Horizontal axis represents time after CNO injection. **(E)** Average changes in NREM, wake and REM following Saline (open circles) or CNO injections (solid circles, n = 9) in *Adcyap1*-Cre mice expressing Gq DREADD in NST (Gq(*Adcyap1*)). Horizontal axis represents time after CNO injection. **(F)** Average changes in NREM, wake and REM state following Saline (open circles) or LPS injections (solid circles, n = 9). Horizontal axis represents time after LPS injection. For D-F, asterisks indicate *p* values for Tukey corrected post-hoc tests where ^∗^*p* < 0.05, ^∗∗^*p* < 0.01, ^∗∗∗^*p* < 0.001. Error bars represent ±SEM.

Fluorescent *in-situ* hybridization (FISH) showed that 63.3 ± 2.4 % (mean ± SEM) of NST^LPS-TRAP^ neurons expressed the glutamatergic marker *Slc17a6* (encoding vesicular glutamate transporter 2) while 36.3 ± 2.9 % expressed the GABAergic/glycinergic marker *Slc32a1* (encoding vesicular GABA transporter) (Fig. 1B). Consistent with a recent study ^24^, we found that the majority of NST^LPS-TRAP^ neurons (56.6 ± 3.8%) expressed *Adcyap1* (encoding pituitary adenylate cyclase-activating polypeptide). Only a small fraction (21.7 ± 4.3 %) expressed *Cartpt* (Fig. S1B-C), suggesting that the sickness-activated neurons are largely distinct from the baroreceptive NST neurons that also promote sleep ^31^.

We then tested the effect of chemogenetic activation of NST^LPS-TRAP^ neurons. Sleep-wake states were measured in freely moving mice in their home cage, based on electroencephalogram (EEG) and electromyogram (EMG) recordings. In mice expressing excitatory DREADD (hM3D(Gq)-mCherry) in NST^LPS-TRAP^ neurons, clozapine-N-oxide CNO (0.3 mg/kg, IP) injection caused a strong increase in NREM sleep and reduction in wakefulness compared to saline control (repeated measures ANOVA; WAKE: *F* _(1,40)_ = 34.0, *p* = 0.0004; NREM: *F* _(1,40)_ = 34.8, *p* = 0.0004, REM: *F* _(1,40)_ = 81.5, *p* = 0.08; Fig. 1D). Chemogenetic activation of *Adcyap1*-expressing NST neurons (NST*^Adcyap1^*) likewise increased NREM sleep while reducing both wakefulness and REM sleep (repeated measures ANOVA; WAKE: *F* _(1,25)_ = 24.2, *p* = 0.004; NREM: *F* _(5,25)_ = 5.1, *p* = 0.002; REM: *F* _(5,25)_ = 5.0, *p* = 0.003; Fig. 1E). Thus, activating NST^LPS-TRAP^ or NST*^Adcyap1^* neurons is sufficient to promote NREM sleep, similar to the effect of LPS injection (repeated measures ANOVA; WAKE: *F* _(1,64)_ = 45.8, *p* = 0.0001; NREM: *F* _(1,64)_ = 100.7, *p* = 0.000008; REM: *F* _(8,64)_ = 4.7, *p* = 0.0001; Fig. 1F).

### NST-to-PB projection promotes sleep

To identify the downstream pathways by which NST sickness-activated neurons promote sleep, we traced their axonal projections by injecting AAV8-pCAG-FLEX-EGFP into the NST of *Adcyap1*-*Cre* mice (Fig. S2A). GFP-labeled axons were observed in multiple brain areas, including the bed nucleus of the stria terminalis (BNST), several hypothalamic and thalamic regions, periaqueductal gray (PAG), amygdala (AMY), LC, PB, and several other brainstem areas (Fig. S2B).

We then selected brain areas with strong NST*^Adcyap1^* projections and known roles in regulating brain states for functional investigation. We expressed a stabilized step function opsin (SSFO; AAV5-EF1a-mW, 5 s/pulse, every 20 ± 5 min) in the PB (Fig. 2A-C), paraventricular thalamus (PVT; Fig. 2D), and ventrolateral periaqueductal gray (vlPAG; Fig. 2E). Activation of the NST→PB projection caused a significant increase in NREM sleep and reduction in wakefulness (repeated measures ANOVA; *F* _(2,30)_ = 30.1, *p* = 0.00006; baseline – laser, wake: *t* _(25)_ = 13.2, *p* = 0.0004; NREM: *t* _(25)_ = -14.6, *p* = 0.001; REM: *t* _(25)_ = 0.5, *p* = 0.6; Fig. 2B,C). In contrast, stimulation of NST→PVT and NST→vlPAG projections caused no significant change in brain states (repeated measures ANOVA; PVT, *F* _(2,24)_ = 0.9, *p* = 0.4; vlPAG, *F* _(2,30)_ = 0.6, *p* = 0.6; Fig. 2C,D).

**Figure 2.**
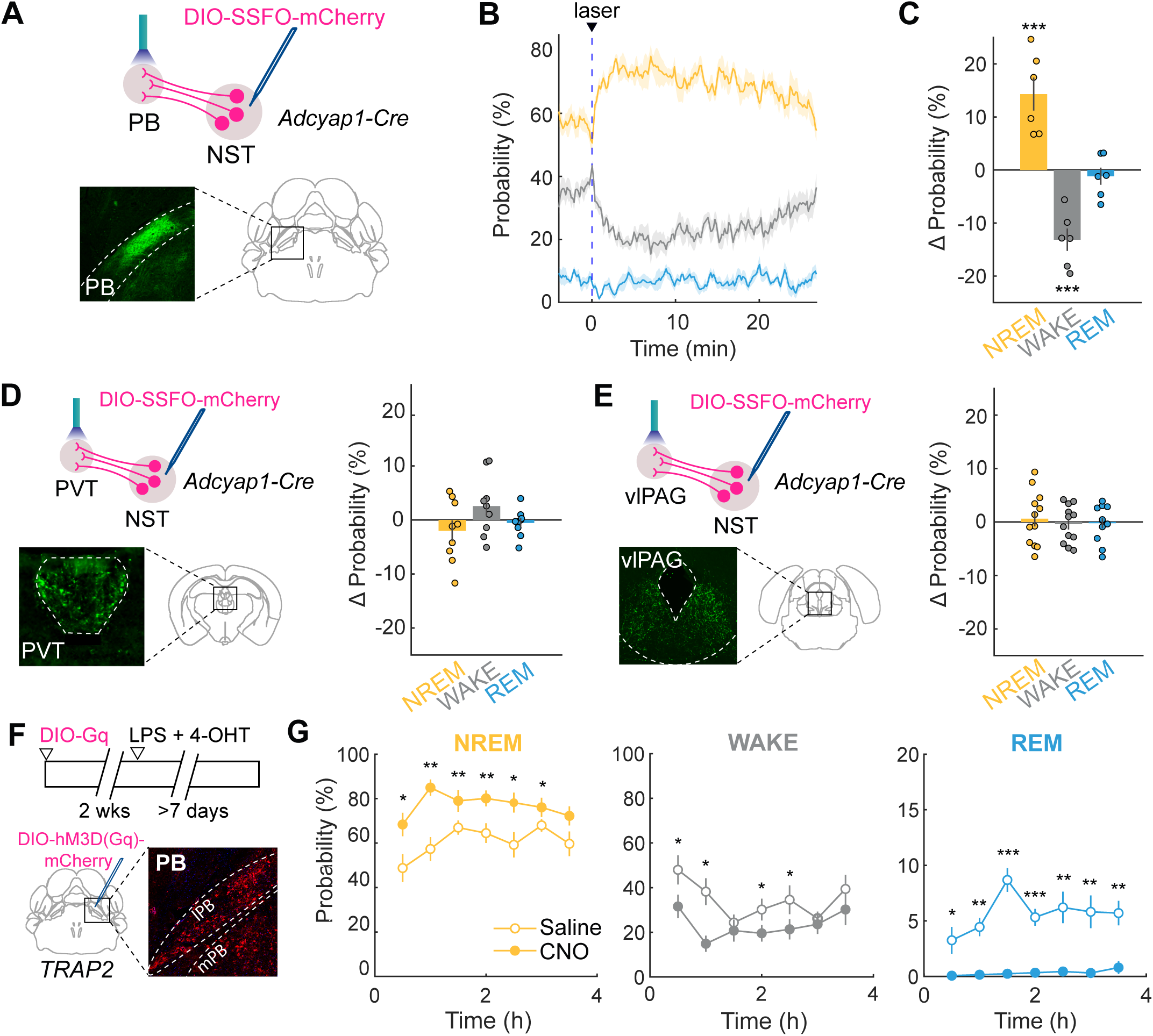
NST-to-PB projection promotes sleep. **(A)** Schematic protocol for optogenetic activation of NST → PB terminals with stabilized step function opsin (SSFO). Optogenetic fibers were placed above PB in *Adcyap1*-Cre mice injected with AAV-DIO-SSFO-mCherry in NST. A 5 s laser pulse was delivered every 20 ± 5 min for each 4-hour session. Inset shows axons from NST*^Adcyap1^* neurons terminating in PB. **(B)** Average laser-evoked change in brain state (NREM, wake, REM) after laser stimulation of NST → PB terminals. Line represents average across individual mice (n = 6), shadow represents ±SEM. Vertical blue line indicates laser onset. **(C)** Quantification of laser-evoked change in brain state in the 20 minutes after laser onset minus 5 min before laser onset. Circles represent individual animals. Error bars represent ±SEM. Asterisks indicate statistical significance where, ^∗^*p* < 0.05, ^∗∗^*p* < 0.01, ^∗∗∗^*p* < 0.001. **(D)** *Left*: Same as (A but for NST→PVT terminals. Inset shows axon terminals from NST*^Adcyap1^*neurons terminating in PVT. *Right:* Same as (A) but for NST→PVT terminal stimulation (n = 9). **(E)** *Left*: Same as a but for NST→vlPAG terminals. Inset shows axon terminals from NST*^Adcyap1^*neurons terminating in vlPAG. *Right:* Same as C but for NST→vlPAG terminal stimulation (n = 12). **(F)** *Top:* Schematic of PB^LPS-TRAP^ experimental protocol. TRAP2 mice were injected with excitatory DREADD (DIO-Gq) in PB. After two wks, mice were co-injected with LPS (0.4 mg/kg) and tamoxifen (4-OHT; 20 mg/kg) to induce Gq DREADD expression in PB^LPS-TRAP^ neurons. *Bottom*: Coronal diagram of PB injection in TRAP2 mice. Inset shows example fluorescence image of PB Gq expression (mCherry). **(G)** Average changes in NREM, wake and REM following Saline (open circles) or CNO injections (solid circles, n = 9) in mice expressing Gq DREADD in PB^LPS-TRAP^ neurons. Horizontal axis represents time after CNO injection. Asterisks indicate *p* values for Tukey corrected post-hoc tests where ^∗^*p* < 0.05, ^∗∗^*p* < 0.01, ^∗∗∗^*p* < 0.001. Error bars represent ±SEM.

To further examine the role of the PB in sickness-induced sleep, we tagged LPS-activated neurons in the PB using TRAP2 mice (Fig. 2F). Chemogenetic activation of PB^LPS-TRAP^ neurons markedly increased NREM sleep and reduced wakefulness and REM sleep (repeated measures ANOVA; WAKE: *F* _(1,35)_ = 21.9, *p* = 0.02; NREM: *F* _(5,35)_ = 50.5, *p* = 0.0002; REM: *F* _(5,35)_ = 2.7, *p* = 0.03; Fig. 2F,G), consistent with the effect of activating NST*^Adcyap1^* → PB projection.

### LPS-induced changes in neuromodulatory systems

Neuromodulators play crucial roles in shaping cognition and emotion ^38–41^, and their dysregulation may contribute to key features of sickness behavior such as anhedonia and fatigue ^42–44^. We thus examined the effects of LPS on three well-known arousal-promoting neuromodulators: norepinephrine (NE), dopamine (DA) and acetylcholine (ACh). GRAB sensor for each neuromodulator ^45–48^ was expressed in several brain regions, and fiber photometry imaging of GRAB fluorescence was used to measure NE, DA or ACh levels (Fig. 3A,E,F). We also injected a Cre-inducible AAV expressing a red-shifted channelrhodopsin (AAV-FLEX-ChrimsonR-tdT) into the LC, ventral tegmental area (VTA), or basal forebrain (BF) of *Dbh-Cre*, *Slc6a3-Cre* or *ChAT-Cre* mice, respectively. This allowed us to measure the release of each neuromodulator evoked by optogenetic activation of the corresponding cell type.

**Figure 3.**
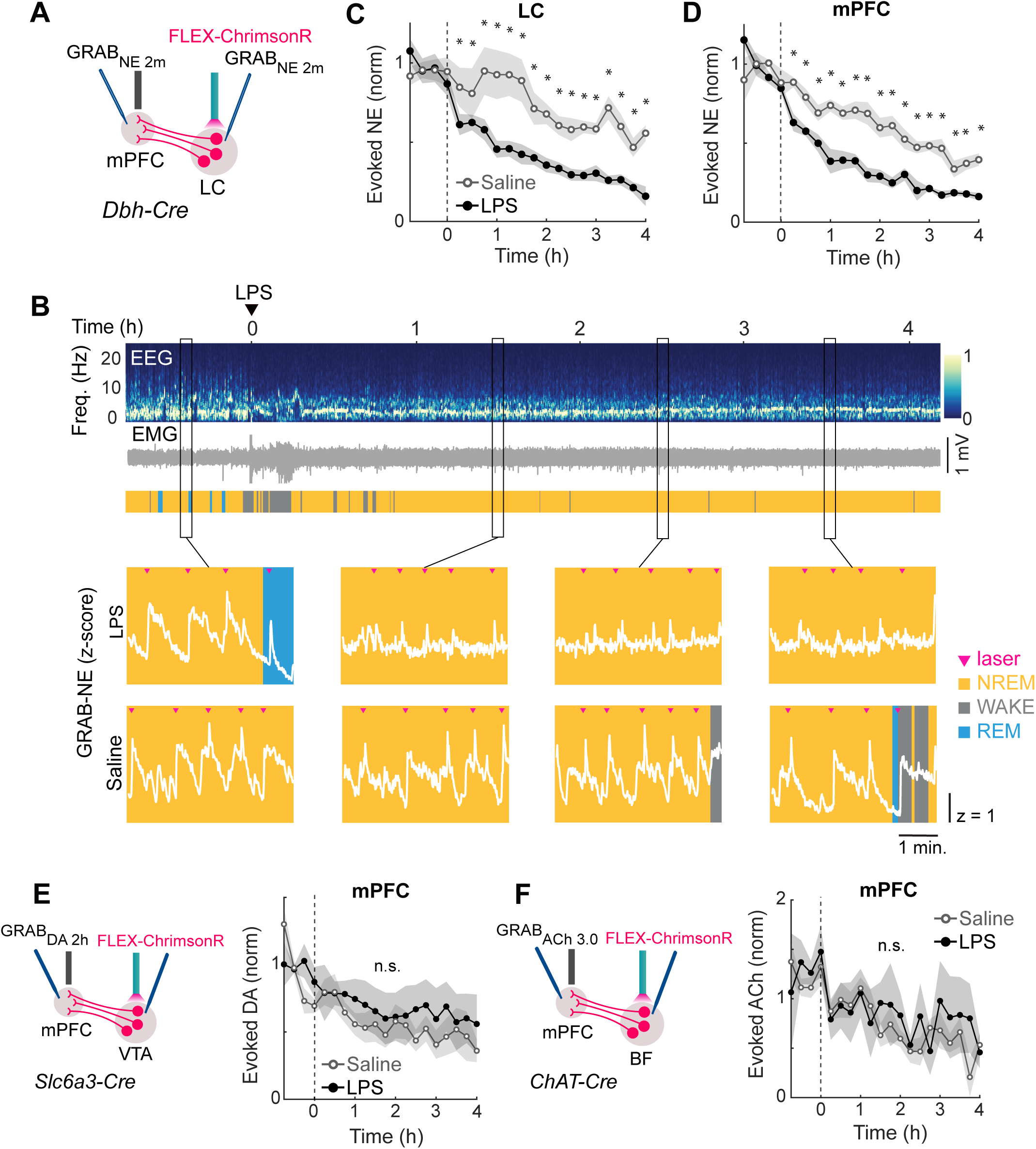
LPS-induced changes in neuromodulatory systems. (A) Schematic for measuring optogenetic laser-evoked release in noradrenergic (NE) neural populations. (B) *Top*: Example LPS session showing EEG, EMG and color-coded brain states. *Bottom:* LC GRAB_NE_ z-scored fluorescence traces overlayed on brain state for selected periods before and after LPS or saline injections. Pink arrowheads indicate laser pulses to evoke release. (C) Laser-evoked GRAB_NE_ response averaged in 15-min bins for LC (n = 12). Horizontal axis indicates time from LPS (black) or Saline (gray) injection. Lines represent average across individual mice. Shadow represents ±SEM. Asterisks indicate *p* values for Tukey corrected post-hoc tests where ^∗^*p* < 0.05. (D) Same as C but for mPFC (n = 7). (E) *Left*: Schematic for measuring optogenetic laser-evoked release in dopaminergic (DA) neural populations. *Right:* Laser-evoked GRAB_DA_ response averaged in 15-min bins for mPFC. Horizontal axis indicates time from LPS (black), or saline (gray) injection. Asterisks indicate *p* values for Tukey corrected post-hoc tests where ^∗^*p* < 0.05. Lines represent average across individual mice (n = 6). Shadow represents ±SEM. (F) *Left:* Schematic for measuring optogenetic laser-evoked release in cholinergic (ACh) neural populations. *Right:* Laser-evoked GRAB_ACh_ response averaged in 15-min bins for mPFC. Horizontal axis indicates time from LPS (black), or saline (gray) injection. Lines represent average across individual mice (n = 10). Asterisks indicate *p* values for Tukey corrected post-hoc tests where ^∗^*p* < 0.05.

We first examined the effect of LPS on NE transmission. Each brief laser pulse in the LC (50 ms/pulse, applied every 60 ± 20 s) evoked a transient increase in GRAB_NE_ fluorescence (Fig. 3B,C), and its amplitude was used to quantify evoked NE release. After LPS injection, evoked NE release in both the LC and mPFC were markedly diminished compared to saline control; this effected started at ∼15 min post-injection and persisted throughout the 4-h recording session (repeated measures ANOVA; LC: *F* _(1,19)_ = 3.3, *p* = 0.00006; mPFC: *F* _(1,19)_ = 2.8, *p* = 0.0002; Fig. 3C,D). We also measured the calcium activity of LC neurons using jGCaMP8s. Optogenetically evoked calcium responses were also strongly reduced following LPS injection (repeated measures ANOVA; *F* _(312,19)_ = 1.87, *p* = 0.02; Fig. S3a), suggesting that the reduction in evoked NE release is at least partly due to reduced excitability of LC neurons.

Next, we measured DA transmission in *Slc6a3-Cre* mice with AAV-FLEX-ChrimsonR-tdT injected into the VTA and AAV9-hSyn-DA2h into both the mPFC and nucleus accumbens (NAc). In contrast to NE release, evoked DA release showed no significant change after LPS injection in the mPFC (repeated measures ANOVA; *F* _(1,19)_ = 2.8, *p* = 0.15; Fig. 3E) and a slight increase in the NAc (repeated measures ANOVA; *F* _(1,19)_ = 29.3, *p* = 0.0000001; Fig. S3B). We also examined evoked ACh release in *ChAT-Cre* mice injected with AAV5-hsyn-ACh3.0 into the mPFC and found no significant change following LPS injection (repeated measures ANOVA; *F* _(1,19)_ = 1.7, *p* = 0.5; Fig. 3F). Taken together, among the three wake-promoting neuromodulators tested, evoked NE transmission was selectively suppressed by LPS injection.

### NST and PB sickness neurons regulate NE transmission

Because activation of NST*^Adcyap1^* neurons was sufficient to re-capitulate the LPS-induced increase in NREM sleep, we next asked whether activating these neurons changes NE release. *Adcyap1-Cre* mice were crossed with *Dbh-Flpo* mice to allow for simultaneous chemogenetic activation of NST*^Adcyap1^* neurons and optogenetic stimulation of LC-NE neurons. We injected Cre-inducible AAV expressing excitatory DREADD (AAV8-hM3D(Gq)-mCherry) into the NST, Flpo-inducible AAV expressing ChrimsonR (AAV8-Ef1a-fDIO-ChrimsonR-tdT) into the LC, and AAV9-hSyn-NE2m into both the LC and mPFC of these mice (Fig. 4A). Chemogenetic activation of NST*^Adcyap1^* neurons with CNO caused a marked reduction of laser-evoked LC-NE release in both the LC and mPFC (repeated measures ANOVA; LC: *F* _(1,19)_ = 2.5, *p* = 0.0005; mPFC: *F* _(1,19)_ = 89.2, *p* = 2×10^-16^; Fig. 4b-d, Fig. S4A), similar to the effect of LPS injection (Fig. 3C,D).

**Figure 4.**
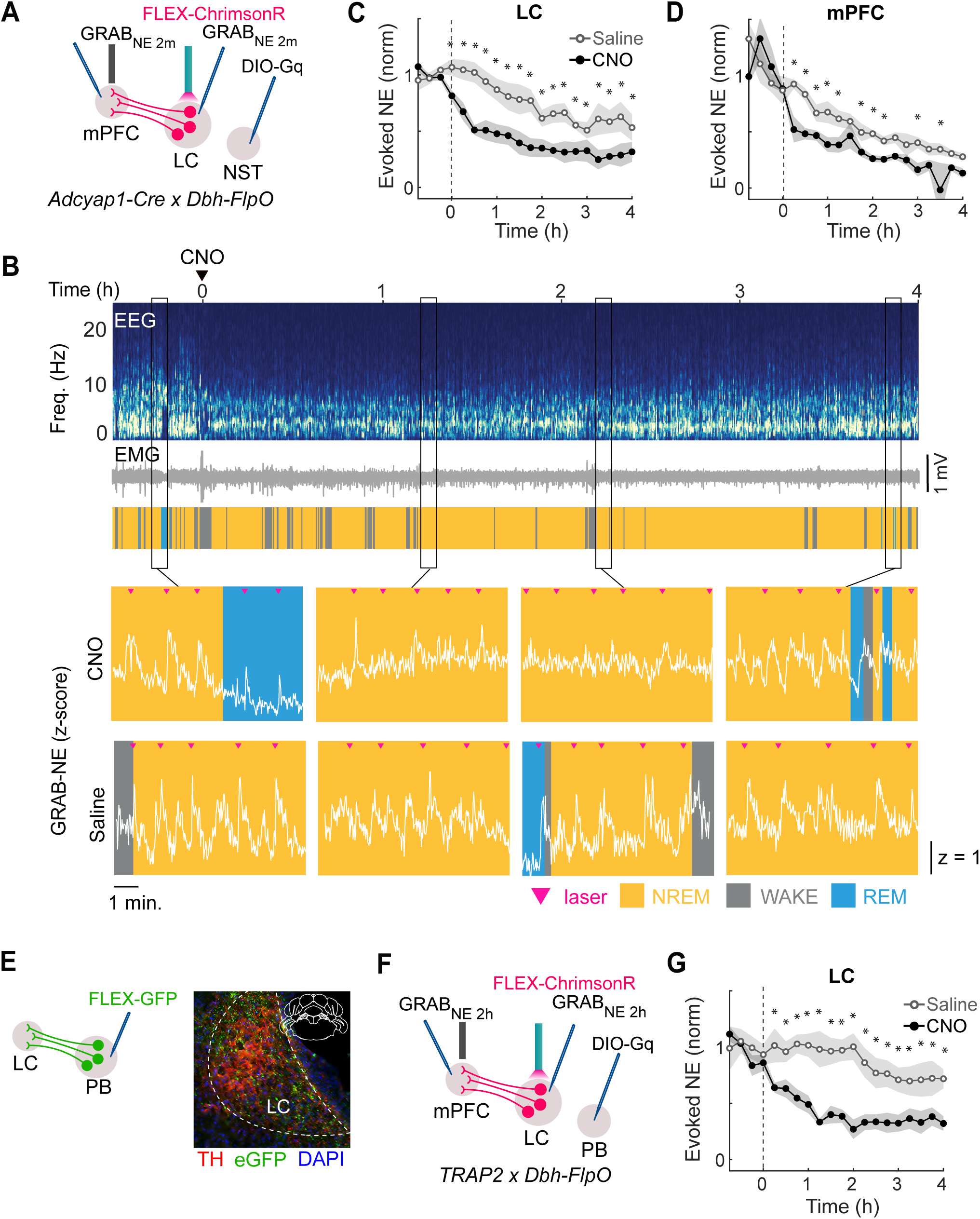
NST and PB sickness neurons regulate NE transmission. (A) Schematic for measuring optogenetic laser-evoked NE release in LC and mPFC during chemogenetic activation of NST*^Adcyap^*neurons. (B) *Top*: Example CNO session showing EEG, EMG and color-coded brain states. *Bottom:* Zoomed LC GRAB_NE_ z-scored fluorescence traces overlayed on brain state for selected epochs before and after CNO (top) or saline (bottom) injections. Pink arrowheads indicate laser pulses to evoke release in LC. (C) Laser-evoked LC-NE responses during chemogenetic activation of NST*^Adcyap^* neurons, averaged in 15-min bins. Horizontal axis indicates time from CNO (black), or saline (gray) injection. Lines represent average across individual animals (n = 9); Shadow represents ±SEM. Asterisks indicate *p* values for Tukey corrected post-hoc tests where ^∗^*p* < 0.05. (D) Same as C but for mPFC LC-NE evoked responses (n = 6). (E) *Top:* Schematic of axonal tracing from PB^LPS-TRAP^ neurons. AAV-FLEX-GFP was injected into the PB of TRAP2 mice to label projections to LC. *Bottom:* Example fluorescence images of LC-NE neurons (TH, red) overlapping with eGFP labeled axons from PB (green). No GFP labelled cell bodies were observed in LC TH+ neurons, indicating virus injections were restricted to the PB target site. (F) Schematic for measuring optogenetic laser-evoked NE release in LC and mPFC during chemogenetic activation of PB^LPS-TRAP^ neurons. (G) Laser-evoked LC-NE responses during chemogenetic activation of PB^LPS-TRAP^ neurons, averaged in 15-min bins. Horizontal axis indicates time from CNO (black), or saline (gray) injection. Lines represent average across individual animals (n =6); Shadow represents ±SEM. Asterisks indicate *p* values for Tukey corrected post-hoc tests where ^∗^*p* < 0.05.

Chemogenetic activation of PB^LPS-TRAP^ neurons also promoted NREM sleep (Fig. 2F,G), and anterograde tracing from these neurons revealed projections to the LC, where GFP-labeled PB^LPS-TRAP^ axons overlapped with tyrosine hydroxylase (TH)-positive LC-NE neurons (Fig. 4E). To test the effect of PB^LPS-TRAP^ neuron activation on evoked NE release, we crossed TRAP2 mice and *Dbh-Flpo* mice and injected AAV8-hM3D(Gq)-mCherry into the PB instead of NST (Fig. 4F). Chemogenetic activation of PB^LPS-TRAP^ neurons markedly reduced evoked LC-NE release (repeated measures ANOVA; *F* _(1,19)_ = 3.3, *p* = 0.00008; Fig. 4G, Fig. S4B), similar to the effect of NST*^Adcyap1^* neuron activation. Thus, activation of either NST or PB sickness neurons reduced evoked LC-NE release.

## DISCUSSION

In this study, we examined the neural pathway originating from the NST that promotes sleep during sickness. Using activity-dependent genetic labeling, we tagged LPS-activated NST neurons and showed that their activation drives an increase in NREM sleep, similar to the effect of LPS injection (Fig. 1). These NST neurons project strongly to the PB, where sickness-activate neurons also promote NREM sleep (Fig. 2). Using GRAB sensors to monitor several wake-promoting neuromodulators (NE, DA, ACh), we showed that NE release evoked by LC stimulation was selectively suppressed by LPS injection (Fig. 3) and by activating NST or PB sickness neurons (Fig. 4).

The NST serves as the primary hub for visceral sensory information entering the brain, integrating and relaying essential signals to regulate autonomic function and maintain homeostasis. Previous studies have shown that LPS-induced sleep increase is significantly diminished by vagotomy or NST inactivation ^23–27^, demonstrating the necessity of this pathway in sickness-induced sleep. Expanding on these findings, we showed that direct activation of NST sickness neurons is sufficient to promote sleep (Fig. 1).

Although NST sickness neurons project to multiple brain regions (Fig. S2), optogenetic activation of their axon terminals showed that only the NST→PB projection strongly promotes NREM sleep (Fig. 2). Like the NST, the PB is a key node in the central autonomic network, regulating a range of behaviors, including feeding, respiration, and thermoregulation ^49^. Previous studies have demonstrated a prominent role of the medial PB in promoting wakefulness ^50–52^. In our study, however, the NST sickness neurons project primarily to the lateral PB (Fig. 2A). In future studies, it would be interesting to determine the molecular identity of PB^LPS-TRAP^ neurons and their interactions with other nodes of the central autonomic network to promote sleep ^53^. Furthermore, while our focus was on circuits that promote sleep, sickness is accompanied by multiple behavioral changes, including loss of motivation, reduced feeding, and autonomic alterations. It will be interesting to determine how the circuits mediating these changes interact with the sleep-promoting pathway we have characterized.

Using genetically encoded biosensors to monitor NE, DA, and ACh levels, we found that peripheral immune activation selectively affects NE transmission. Evoked NE release was markedly reduced following either LPS injection or activation of NST and PB sickness neurons (Figs. 3, 4), suggesting that these circuits promote sleep in part through diminished NE transmission. Notably, unlike evoked NE release, the overall NE level was elevated by activation of NST sickness neurons (Fig. S4). This is consistent with previous studies based on microdialysis ^54–57^, and it may be part of a negative feedback mechanism that helps to dampen inflammation ^58–61^. While the opposing changes in overall and evoked NE levels may appear contradictory in their effects on sleep, accumulating evidence suggests that transient NE increases promote arousal, whereas sustained NE elevation may in fact enhance NREM sleep ^62–65^.

In summary, we have identified an NST→PB pathway that mediates sickness-induced sleep, in part by modulating LC-NE transmission. Besides peripheral immune activation, LC-NE signaling is also regulated by microglia, the brain’s resident immune cells ^66^. As a potent wake-promoting neuromodulator regulated by both neuronal and non-neuronal mechanisms, the NE system is well positioned to link the body’s need for recovery from sickness to the homeostatic regulation of sleep drive.

## METHODS

### Animals

All procedures were performed in accordance with the protocol approved by the Animal Care and Use Committee at the University of California, Berkeley. Adult (6-12 weeks old) male and female mice were used for all experiments. Mice were kept on a 12:12 light:dark cycle (lights on at 07:00 am and off at 07:00 pm) with free access to food and water. After virus injections and surgical implantation of EEG/EMG electrodes and optical fibers, mice were individually housed to prevent damage to the implant before experiments. Experiments were conducted at least 2 weeks after surgery. Mice utilized here include: TRAP2: Jackson strain # 030323; *Adcyap1*-Cre: Jackson strain # 030155; Dbh-Cre: B6.FVB(Cg)-Tg(Dbh-Cre)KH212Gsat/Mmucd, MMRRC: 036778-UCD; Dbh-Flpo: Jackson strain # 033952; *Slc6a3*-Cre: Jackson strain # 006660; CHAT-Cre: Jackson strain # 06410.

### LPS administration

Lipopolysaccharide from Escherichia coli (O111:B4, Sigma L2630) was reconstituted in saline (1 mg/ml^−1^) and frozen into single-use aliquots. Further dilutions in saline to the experimental dose (0.4 mg/kg) were prepared before each experiment. All doses were delivered by intraperitoneal injection.

### TRAP induction

4-hydroxytamoxifen (4-OHT) was prepared based on K. Deisseroth’s lab protocol. For LPS-TRAP, 4-OHT (2 mg/ml in saline with 2% Tween-80, 20 mg/kg) and LPS (0.4 mg/kg) were co-injected intraperitoneally. Mice were given at least 7 days to allow for expression before beginning experiments. To measure the overlap between LPS-TRAP and LPS-Fos, mice were sacrificed and perfused 2 hours after LPS injections.

### Surgical procedures

Adult mice (6 - 12 weeks old; male and female) were anesthetized with isoflurane (5% induction, 1.5% maintenance) and placed on a stereotaxic frame. Buprenorphine (0.1 mg/kg, subcutaneous) and meloxicam (10 mg/kg subcutaneous) were injected before surgery. Lidocaine (0.5%, 0.1 mL, subcutaneous) was injected near the target incision site. Body temperature was stably maintained throughout the procedure using a heating pad. After asepsis, the skin was incised to expose the skull and overlying connective tissue was removed.

For EEG and EMG recordings, a reference screw was inserted into the skull on top of the left cerebellum. EEG recordings were made from two screws on top of the left and right cortex at -3.5 (AP) ± 3.5 mm (ML). Two EMG electrodes were inserted into the neck musculature. Insulated leads from the EEG and EMG electrodes were soldered to a pin header, which was secured to the skull using dental cement.

Virus injections were performed as above but a craniotomy was made on top of target regions (see below for coordinates) and 50-200 nanoliters of virus was injected using a Nanoject II (Drummond) and a glass micropipette. For optogenetic and fiber photometry experiments, fiber optic ferrules (1.25 mm ferrule, 200um Core, 0.39NA) were stereotactically inserted and secured to the skull using dental cement. All experiments were performed at least two weeks after surgery to allow for virus expression and animal recovery.

The following stereotaxic coordinates were used for virus injections and optogenetic cannula placement. Unless otherwise noted, coordinates are listed relative to Bregma.

NST: -7.3 AP, 0.25 ML, -5.0 DV

PVT: -0.4 AP, 0 ML, -3.6 DV

vlPAG: -4.9 AP, 0.6 ML, -2.7 DV

NAc: + 1.4 AP, 1.2 ML, - 4.0 DV from dura

VTA: -3.2, 0.5, 4.1 from dura

mPFC: +2.1 AP, 0.3, -1.6 DV

BF: +0.1 AP, 1.5 ML, 5.3 DV

LC: -5.5 AP, 0.9 ML, -3.8 to -3.2 DV. For LC 50 nL were injected every 0.2 mm at multiple depths.

Ferrules were placed at -3.65 DV.

PB: -4.9 AP, 1.5 ML, -3.7 DV

### Viruses

AAV5-hSyn-DIO-hM3D(Gq)-mCherry (# 44361), AAV8-pCAG-FLEX-EGFP-WPRE (# 51502) and AAV5-hSyn-FLEX-ChrimsonR-tdT (# 62723) were obtained from Addgene. AAV5-EF1a-DIO-SSFO-mCherry was obtained from the University of North Carolina (UNC) vector core. Grab sensors AAV9-hSyn-NE2m, AAV5-hsyn-ACh3.0(ACh4.3), and AAV9-hSyn-DA2h(DA4.3), were obtained from WZ Biosciences.

### Polysomnographic recordings

Behavioral experiments were carried out in home cages placed in sound-attenuating boxes between 9:00 am and 7:00 pm. EEG/EMG electrodes were connected to flexible recording cables via a mini-connector. EEG/EMG signals were acquired using a TDT PZ5 amplifier and Synapse software, with a bandpass filter of 0.3–500 Hz and sampling rate at 1017 Hz. Spectral analysis was carried out using fast Fourier transform, and brain states were classified as described previously (wake: desynchronized EEG and high EMG activity; NREM: synchronized EEG with high-amplitude, low-frequency (1–4 Hz) activity and low EMG activity; REM: high EEG power at theta frequencies (6–9 Hz) and low EMG activity). The classification was determined using 5 second bins and with a custom-written graphical user interface (programmed in MATLAB, MathWorks).

### Fiber photometry recording

Fiber photometry recording was performed using TDT RZ10x real-time processor. Fluorescence elicited by 405 nm and 465 nm LEDs were filtered through the dichroic mini cube (Doric lenses) and collected with an integrated photosensor on the RZ10x. Signals were demodulated and pre-processed using the TDT Synapse software collected at a sampling frequency of 1017 Hz.

For LC evoked stimulation experiments, a patch able from the 635 red laser diode (RWD Life Science) were connected to the dichroic mini-cube (Doric) to enable simultaneous optogenetic laser stimulation and fiber photometry from the same fiber tip.

### Immunohistochemistry and fluorescence in situ hybridization (FISH)

Mice were deeply anaesthetized and trans-cardially perfused with 0.1M PBS followed by 4% paraformaldehyde in PBS. For fixation, samples were kept overnight in 4% paraformaldehyde. Samples were then placed in a 30% sucrose solution for 24-48 h for cryoprotection. After embedding and freezing, brains were sectioned into 30µm (FISH samples) or 50 µm (for other immunohistochemistry) coronal slices. For immunohistochemistry, brain slices were washed using PBS three times, permeabilized using PBST (0.3% Triton X-100 in PBS) for 30 min and then incubated with blocking solution (5% normal goat serum or normal donkey serum in PBST) for 1 hr followed by primary antibody incubation overnight at 4° C. Antigen retrieval pretreatment was performed prior to c-fos antibody treatment. The next day, slices were washed with PBS and incubated with appropriate secondary antibodies for 2 h at room temperature. FISH was performed using RNAscope Multiplex Fluorescent Assays V2 according to the manufacturer’s instructions (Advanced Cell Diagnostics). Fluorescence images were taken using a fluorescence microscope (Keyence BZ-X710) or a high throughput slide scanner (Nanozoomer-2.0RS, Hamamatsu).

### Quantification and statistical analysis

For fiber photometry experiments, the 405-nm channel was used to correct nonspecific, calcium-independent changes in fluorescence, e.g., movement artifacts. Each channel (465-nm and 405-nm) was first fit with a single exponential to remove the baseline change due to bleaching. The 405-nm signal was then fit to the 465-nm signal using a least-squares linear fit method^67^ and then subtracted from the 465-nm signal. The resulting signals were then converted to z-scores based on the mean and standard deviation of the entire imaging session.

For changes in overall level, corrected photometry signals for each session type (saline, CNO) were averaged into 15-min bins and subtracted for each mouse (CNO - saline). This difference was then averaged across mice to assess changes in overall level (Fig. S3, Fig. S4). For laser-evoked release, 465 channel signals were z-scored and normalized to the mean of the first hour of the 5-hour session. Evoked response for each pulse was calculated as the difference between the time period 1s after laser onset and the 1 sec before laser onset. Evoked responses were normalized to the first hour of the recording session and binned every 15 min. To control for differences in evoked release across brain states, we restricted all analyses to periods of NREM sleep.

Statistical analyses were conducted using MATLAB and R. Most analyses utilized two-way repeated measures mixed ANOVAs implemented in R. We specified random slopes and intercepts models and included mouse/subject as a random covariate using the lme4 package. Treatment (CNO, LPS, or saline) and time (binned by hours or minutes of the recording session) were included as fixed effects. Following a significant main effect or interaction, we conducted post hoc analyses between groups across time. Reported post hoc analyses are *t* tests with Tukey corrections for multiple comparisons and were conducted using the lsmeans package in R. All statistical comparisons are conducted on animal averages (i.e., each animal has one observation per level(s) of the independent variable).

## ACKNOWLEDGEMENTS

We thank Hongfeng Gao for administrative support. Yiyan Hao and Dillon Leung assisted in early phases of the project. We are grateful to members of the Dan lab for helpful discussions.

## AUTHOR CONTRIBUTIONS

Conceptualization, DD and YD; Methodology, DD and YD; Conducting experiments, DD, YY, JS, CC; Data analysis, DD, JS, DS; Writing – Original Draft, DD and YD; Writing – Review and Editing, DD, YD; Supervision, YD; Project Administration, YD; Funding Acquisition, YD

## RESOURCE AVAILABILITY

### Lead Contact

Further information and requests for resources and reagents should be directed to and will be fulfilled by the lead contact, Yang Dan

### Materials availability

This study did not generate new unique reagents.

### Data and code availability

Any additional information required to reanalyze the data reported in this paper is available from the lead contact upon reasonable request.

---

**Figure S1.**
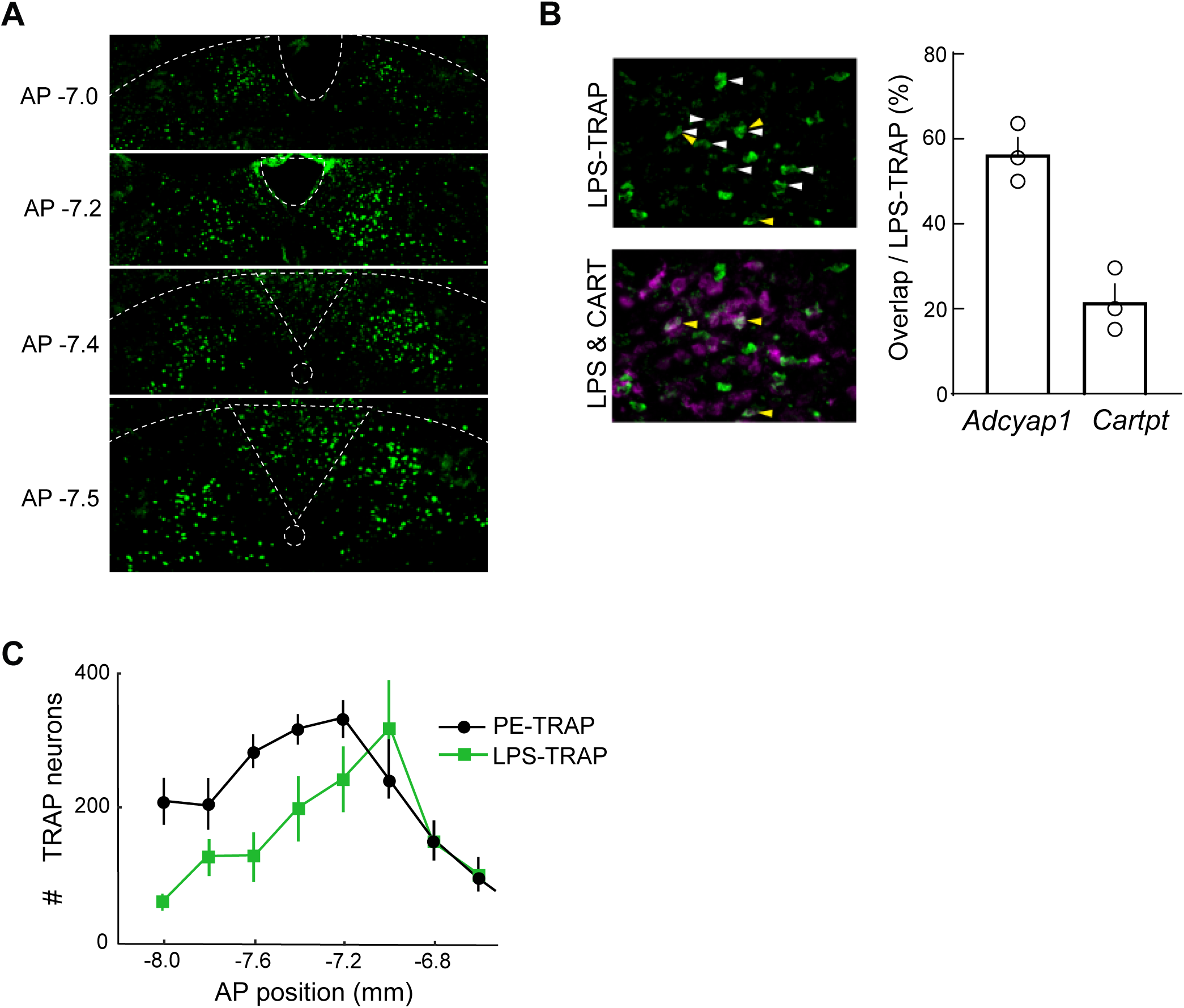
NST^LPS-TRAP^ neurons are distinct from baroreceptive NST neurons. **(A)** Example fluorescence images of *eGFP* labeled NST^LPS-TRAP^ neurons across different anteroposterior planes. **(B)** *Left*: Overlap between NST^LPS-TRAP^ neurons with *Cartpt* marker shown by double FISH. *Right*: Percent overlap between NST^LPS-TRAP^ neurons and *Cartpt* or *Adcyap1* markers. Individual samples are shown as open circles. **(C)** NST^LPS-TRAP^ and baroreceptive NST^PE-TRAP^ ^31^ neuron counts across anteroposterior planes of NST.

**Figure S2.**
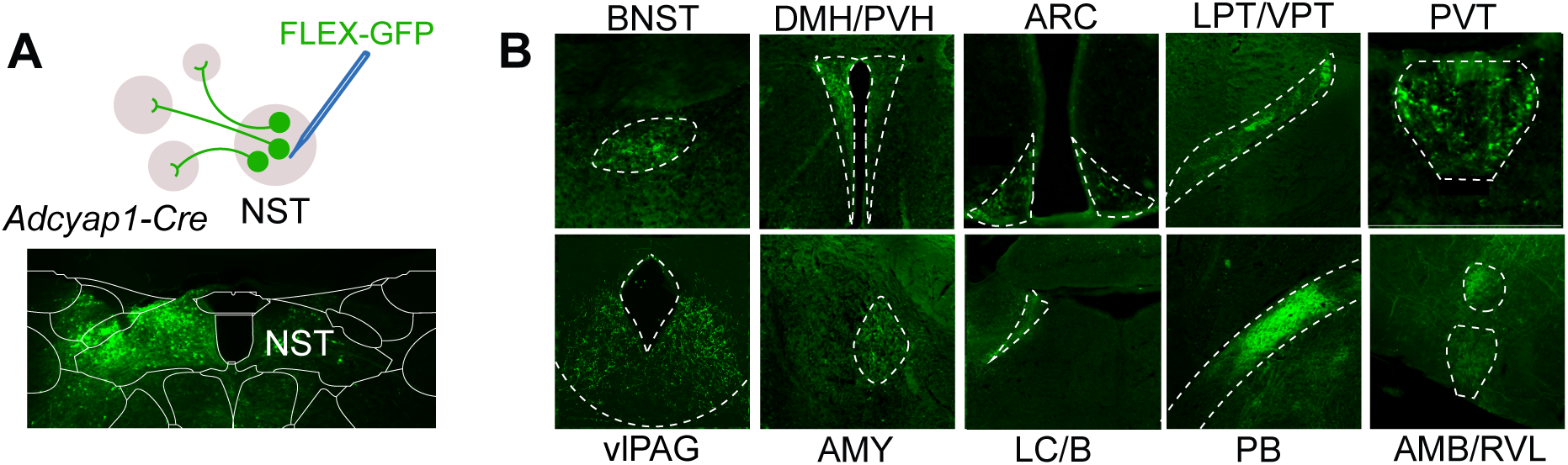
Projections of NST*^Adcyap^* neurons. **(A)** *Top:* Schematic of axonal tracing from NST*^Adcyap^* neurons. AAV-FLEX-GFP was injected into the NST of *Adcyap1*-Cre mice to label axons to projection targets. *Bottom:* Example fluorescence image of GFP labeled NST*^Adcyap^*neurons with atlas overlay. **(B)** Identified projection targets of NST*^Adcyap^* neurons. GFP labeled axons were identified in 14 brain areas including the bed nucleus of the stria terminalis (BNST), dorsomedial and paraventricular hypothalamus (DMH/PVH), arcuate nucleus of the hypothalamus (ARC), lateral posterior and ventroposterior thalamus (LPT/VPT), periventricular thalamus (PVT), periaqueductal gray (PAG), amygdala (AMY), locus coeruleus and Barrington’s nucleus (LC/B), parabrachial (PB), nucleus ambiguous and the rostral ventrolateral medulla (AMB/RVLM).

**Figure S3.**
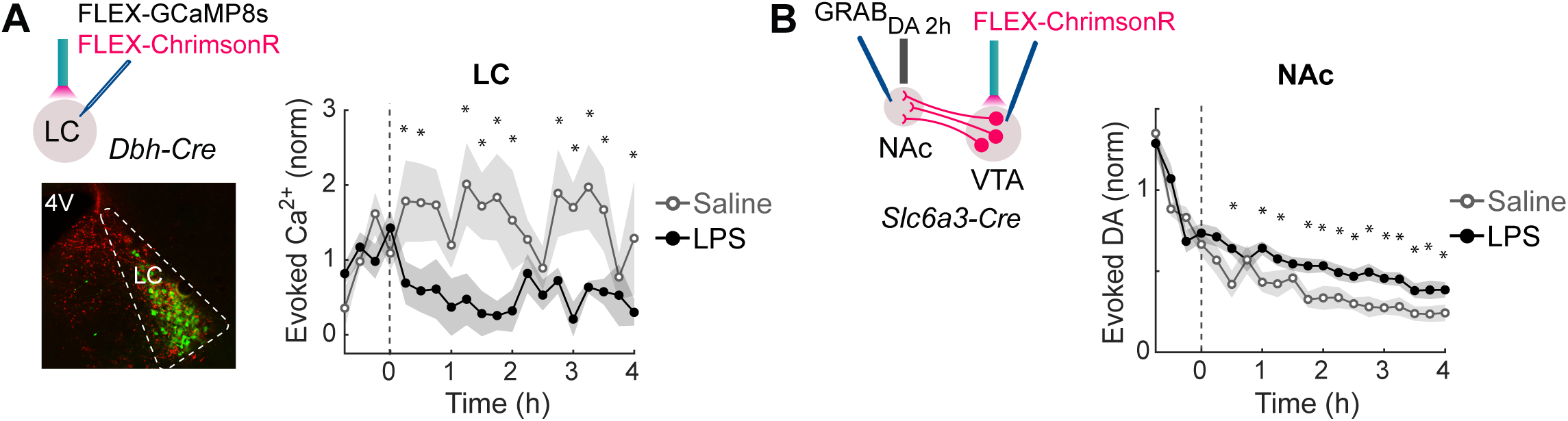
LPS-induced changes in LC evoked calcium response and DA release in NAc evoked by VTA stimulation. **(A)** *Top left:* Schematic for measuring optogenetic laser-evoked calcium responses in LC. *Bottom left*: Example fluorescence image of LC-NE neurons expressing ChrimsonR (tdTomato) and jGCaMP8s. *Right:* Laser-evoked calcium responses averaged in 15-min bins for LC (n = 9). Horizontal axis indicates time from LPS (black) or Saline (gray) injection. Lines represent average across individual mice. Shadow represents ±SEM. Asterisks indicate *p* values for Tukey corrected post-hoc tests where ^∗^*p* < 0.05. **(B)** *Left:* Schematic for measuring optogenetic laser-evoked release in VTA→NAc dopaminergic (DA) neural populations. *Right:* Laser-evoked DA response averaged in 15-min bins for NAc. Horizontal axis indicates time from LPS (black), or saline (gray) injection. Asterisks indicate *p* values for Tukey corrected post-hoc tests where ^∗^*p* < 0.05. Lines represent average across individual mice (n = 12). Shadow represents ±SEM.

**Figure S4.**
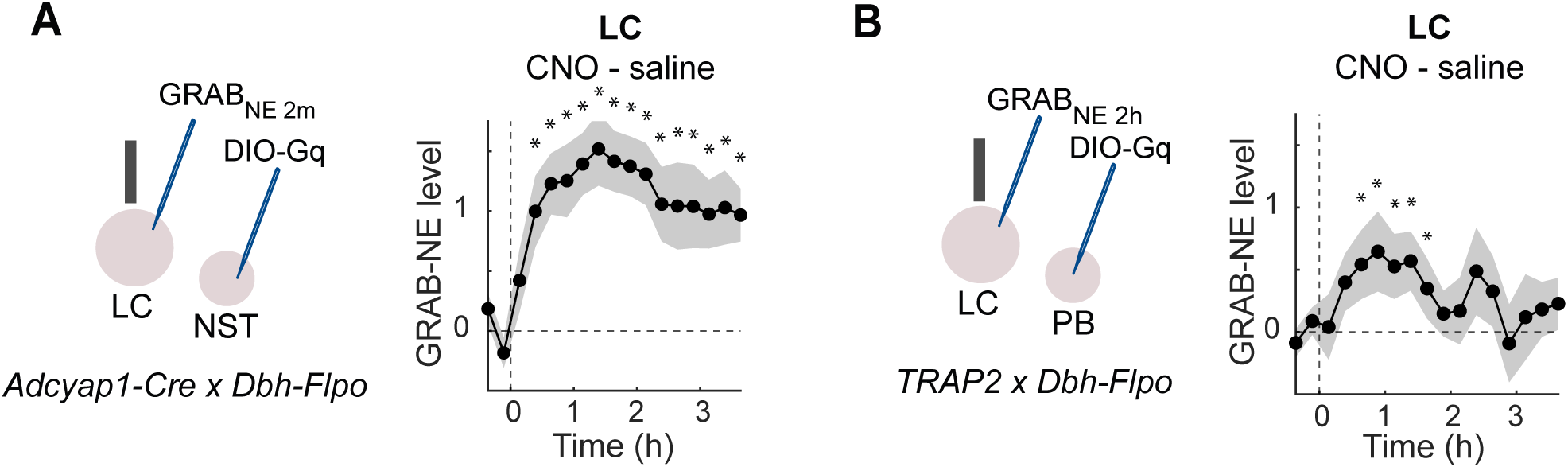
Changes in overall NE level induced by chemogenetic activation of NST or PB sickness neurons. **(A)** *Left*: Schematic for measuring LC NE level during chemogenetic activation of NST*^Adcyap^*neurons. *Right*: Average difference (CNO - Saline; n = 9) of LC GRAB_NE_ z-scored fluorescence traces averaged into 15-min bins during chemogenetic activation of NST*^Adcyap^* neurons. Horizontal axis indicates time from injection. Shadow represents ±SEM. Asterisks indicate *p* values for Tukey corrected post-hoc tests where ^∗^*p* < 0.05. **(B)** *Left*: Schematic for measuring LC NE level during chemogenetic activation of PB^LPS-TRAP^ neurons. *Right*: Average difference (CNO - Saline; n = 6) of LC GRAB_NE_ z-scored fluorescence traces averaged into 15-min bins during chemogenetic activation of PB^LPS-TRAP^ neurons. Horizontal axis indicates time from injection. Shadow represents ±SEM. Asterisks indicate *p* values for Tukey corrected post-hoc tests where ^∗^*p* < 0.05.

